# Microbiota and Small Cell Lung Cancer: A casual bystander or a hidden culprit?

**DOI:** 10.1101/2025.09.26.678903

**Authors:** I Rolim, A Lopez-Beltran, M Pantarotto, E de Sousa, J Sobral, C Farver, N Gil, C Penha-Gonçalves

**Author notes:** **Corresponding Author:** Ines Rolim.

## Abstract

The tumor-associated microbiome is a key player in cancer development, progression, prognosis, and therapeutic response. Notably, distinct microbial signatures have been identified across cancer types. Small cell lung carcinoma (SCLC) accounts for approximately 15% of all lung cancer, yet its microbiome remains unclear. Analyzing the bacteriome composition in tissue from ten SCLC cases and in 10 cases of a heterogenous lung pathology group, we found a distinct microbial signature associated with SCLC with significantly lower diversity and higher dissimilarity, characterized by a higher relative abundance of Firmicutes and Bacteroidota, and a markedly different set of dominant genera (*Pseudomonas, Streptococcus* and *Haemophilus*) resulting in an increased Proteobacteria-to-Actinobacteria ratio. Unexpectedly, mycobiome analysis comparing pooled samples of these SCLC cases with ten pooled lung adenocarcinoma (LUAD) cases revealed that the fungal genus *Taphrina* was uniquely represented in SCLC. Strikingly, mycobiome individual analysis of twenty-one additional SCLC cases compared with 10 LUAD cases showed an increased prevalence of *Taphrina* sp. in SLCL tissue. Overall, the results suggest that SCLC microbiome is distinct from other lung pathologies and uncovers a novel link between the biotrophic plant pathogenic *Taphrina* and human cancer.

## INTRODUCTION

For more than a century the links between cancer and microbes, such as bacteria and viruses, have been explored (1). Recent studies have highlighted the tumor-associated microbiome as a novel player in cancer development, progression, prognosis, and therapeutic response (2–5). Locally resident microbiota, in particular bacteria, have been associated with increased susceptibility to various cancers, including oral squamous cell carcinoma, pancreatic cancer, lung cancer and breast cancer, among others (6). This can result from direct synthesis of carcinogenic compounds exerting genotoxic effects, direct inactivation of genes, initiation of local inflammatory processes leading to alterations in the local microenvironment, or impairment of the immune system’s functionality, resulting in immunosuppression (7). The healthy human respiratory tract is inhabited by diverse and dynamic microbial communities with a crucial role in host protection by regulating innate and adaptive lung immunity since early life (8–11). Yet, the function of lung mycobiota in the maintenance of lung homeostasis has remained largely unknown (12). Microbial organisms have long been recognized as causative agents of infectious diseases and contributors to human morbidity. However, growing evidence now supports their involvement in the etiopathogenesis of non-infectious conditions, including cancer. Several studies have demonstrated that imbalances in the lung microbiota are associated with the development of major chronic respiratory diseases, including chronic obstructive pulmonary disease (COPD), idiopathic pulmonary fibrosis (IPF), asthma, and lung cancer (13-15).

Among all human-infecting microorganisms, fungi remain one of the least studied and understood, despite their substantial impact on human health. Fungal infections affect billions of people worldwide, contributing to over 2 million deaths annually (16,17). While fungi have been implicated in tumorigenesis, evidence is still scarce and their precise role and mechanisms in cancer development remain largely unknown (18,19). *Alternaria arborescens*, a plant-pathogenic species, was recently identified as the most relevant fungus in non-small cell lung carcinoma. Its spores, one of the most effective air allergen, can invade human respiratory tracts and cause respiratory and lung diseases (20). To date, no data on small cell lung carcinoma (SCLC) microbiome has been reported.

SCLC has historically been considered a molecularly homogeneous and highly recalcitrant tumor, with treatment strategies proving insufficient to overcome its aggressive nature (4). New insights into the development and progress of this intriguing neoplasia are needed.

In this exploratory study, a distinct group of 9 SCLC cases emerged among the 80 lung specimens examined (50 neoplastic and 30 non-neoplastic/benign). These specimens were sequenced for the V4 hypervariable region of the 16S ribosomal RNA gene, which informed the subsequent study strategy, including sequencing of the ITS region for fungal identification. We anticipate that this first report on the SCLC microbiome will encourage further investigations into microbial profiles, particularly the mycobiome community, to shed light on the pathogenesis of this aggressive neoplasm.

## MATERIALS AND METHODS

### Ethical permit

This study was approved by the Ethics Committee of Champalimaud Foundation as part of the project *The Pulmonary Microbiome in Lung Carcinoma*, following the European and Portuguese clinical research regulations.

### Case selection

The pathology database of Champalimaud Clinical Centre in Lisbon, Portugal, was searched for biopsies/ surgical specimens with the diagnosis of eight distinct pulmonary pathologies: lung adenocarcinoma (LUAD), lung squamous cell carcinoma (LUSC), typical carcinoid tumor (TCT), small cell lung cancer (SCLC), adenocarcinoma in situ (AIS), pulmonary hamartoma (PH), emphysema, and granulomatous inflammation, accessed between 2018 and 2020. Ten cases per category were selected. A second search was conducted to identify additional cases of SCLC and LUAD. Twenty-one new cases of SCLC and ten cases of LUAD, from 2021 and 2023, were selected for inclusion in the study. The entire collection was systematically analyzed and the diagnosis confirmed by two pathologists (I.R. and A.L.B.), with an average of five (range: 1-8) stained slides evaluated per case, of routinely fixed in 10% neutral buffered formalin and embedded in paraffin (FFPE) material, including H&E and immunohistochemical stains.

### Collection characterization

A total of 111 cases, corresponding to 60 (54%) bronchial/transbronchial/ transthoracic biopsies and 51 (46%) surgical specimens were selected. Of these, 31 cases were SCLC specimens, including 29 biopsies and two surgical specimens. 16S rRNA gene sequencing was performed using DNA isolated from FFPE samples and five paraffin samples from the empty margins of randomly selected specimens were used to control for embedding material background,

### DNA extraction

Five serial 10-μm thick sections were cut from each formalin-fixed, paraffin-embedded (FFPE) sample. The DNA extraction was performed following the protocol from the DNA Sample Preparation Kit (cobas®), with the incubation with Proteinase K extended overnight. Five paraffin samples, consisting of tissue-free paraffin collected from the periphery of the randomly selected specimens, were included as control of the embedding preservation.

### Targeted metagenomic sequencing and taxonomic assignment

To assess microbial community composition, we targeted two distinct genetic regions: the V4 hypervariable region of the 16S ribosomal RNA (rRNA) gene for bacterial identification and the internal transcribed spacer (ITS) region for fungal identification. Protocols and primer pairs recommended by the Earth Microbiome Project were used for both regions. The V4 region of the 16S rRNA gene was amplified with the primer pair 515F (5′-GTGCCAGCMGCCGCGGTAA-3′) and 806R (5′-GGACTACHVGGGTWTCTAAT-3′) (21–23). For the fungal community, the ITS1-ITS2 region was amplified using the ITS1f (5′-CTTGGTCATTTAGAGGAAGTAA-3′) and ITS2 (5′-GCTGCGTTCTTCATCGATGC-3′) primers, as outlined by Smith and Peay (24). Library preparation and amplicon sequencing (Illumina Miseq, 2x 250 cycles) were performed in the Genomics Facility at the Instituto Gulbenkian de Ciência. Primers were removed from high-throughput sequencing reads using Cutadapt (25). Sample inference and fungal taxonomic assignment was performed using DADA2 (26). UNITE database version 9 was used as reference (27).

### Semi-nested PCR and Sanger Sequencing

The universal fungal ITS region (ITS1-ITS2) was targeted for evaluation by semi-nested PCR. For the first PCR, the primer sequences used were ITS1 (5’-TCCGTAGGTGAACCTGCGG-3’) and ITS4 (5’-TCCTCCGCTTATTGATATGC-3’). The 25 µL of reaction mixture contained 30 µg of genomic DNA, the DreamTaq Green PCR Master Mix (Thermo Scientific™, Waltham, MA, USA) and the primers ITS1 and ITS4 (10 µM). The expected band size was around 480 bp. In the second PCR step, the ITS1 and ITS2 (5’GCTGCGTTCTTCATCGATGC-3’) regions were amplified using 1 µl of diluted (1/10) product of the first PCR. So, the product of the first PCR was run into the second PCR to amplify a 170-300 bp DNA, using the same volume of the reaction mixture applied in the first PCR. PCR was carried out under the following conditions: preliminary denaturation at 95 °C—3 min; further for 35 cycles: 95 °C denaturation—l30 s, 56°C annealing for the first PCR and 58.5ºC for the second PCR—30 s, 72 °C synthesis—1 min; final cycle 72 °C—5 min. An aliquot of Taphrina Deformans DNA was used as a positive control. We ran the PCR reaction on the C1000 Touch™ Thermal Cycler (BioRad). The analysis of the PCR products was performed in QIAxcel Advanced (Qiagen). The primer sequences were obtained from Ashraf et al. (28).

### Sanger Sequencing

Briefly, the PCR product of all fungal ITS region amplifications (except sample 2S5) was cleaned using the QIAquick PCR Purification Kit (Qiagen) and sent to sequencing using the ITS1 and ITS2 primers. The PCR product of sample 2S5 was run on a 3% agarose gel to separate the two bands of 222 bp and 251 bp. Both bands were excised and extracted using the QIAquick Gel Extraction Kit (Qiagen). The purified PCR products were sequenced using the Sanger method with BigDye Terminator v3.1 chemistry on an ABI 3730xl DNA Analyzer, performed by STAB VIDA (Portugal). High-quality sequences were aligned and identified using BLAST against the NCBI nucleotide database (https://blast.ncbi.nlm.nih.gov/Blast.cgi). Eligibility was determined based on a high-confidence BLAST match (≥97% identity, E-value < 1e-20) and a minimum read support of 10 reads.

### Statistical analysis

For microbiome analysis (16S rRNA sequencing), alpha diversity (diversity of species within each group) was assessed using the Shannon-Wiener index (H’), Effective Number of Species (ENS), Gini-Simpson (1-D) and the Pielou evenness (J’) indices; beta diversity (diversity of species between groups) was assessed using Principal Coordinate Analysis (PCoA) based on a Bray-curtis dissimilarity matrix, PERMANOVA test and the Jaccard similarity coeficient. Groups were compared using the Kruskal-Wallis Test. Statistical significance was set at p < 0.05. Data statistics and bioinformatic analyses were performed using DATAtab software package (DATAtab Team (2024). DATAtab: Online Statistics Calculator. DATAtab e.U. Graz, Austria. URL https://datatab.net), and R 4.3.1 (R Core Team, 2023), https://www.R-project.org/.

## RESULTS

### Study workflow

A total of 80 lung specimens representing eight distinct pulmonary pathologies were included: lung adenocarcinoma (LUAD), lung squamous cell carcinoma (LUSC), typical carcinoid tumor (TCT), small cell lung cancer (SCLC), adenocarcinoma in situ (AIS), pulmonary hamartoma (PH), emphysema, and granulomatous inflammation - 10 cases per category. All samples underwent sequencing of the V4 hypervariable region of the 16S ribosomal RNA (rRNA) gene for bacterial profiling. The SCLC group exhibited eligible results in 9 of the 10 cases, corresponding to a total of 12105 number of Amplicon Sequence Variants (ASVs), was therefore designated as one of the two study cohorts (1^st^ SCLC cohort). A control group (GC1), with eligible results corresponding to a total of 1673 number of Amplicon Sequence Variants (ASVs) was created, comprising 9 individual cases - TCT, AIS, PH (n=2), emphysema (n=4), and granulomatous inflammation.

Mycobiome analysis was subsequently conducted using Illumina sequencing of the internal transcribed spacer (ITS) region for fungal identification in two pooled groups – 1^st^ SCLC cohort (n=10) and a LUAD group (n=10), designated control group 2 (CG2). A second search was conducted to identify additional cases of SCLC and LUAD. A total of 31 additional cases, corresponding to twenty-one SCLC and ten LUAD, from 2021 and 2023, were selected to enter the study, forming the 2^nd^ SCLC cohort and the third control group (CG3), respectively. A second approach methodology for mycobiome analysis was followed.

Semi-nested PCR targeting the universal fungal ITS region (ITS1-ITS2) for the first PCR and ITS1 and ITS2 regions for the second PCR step, followed by Sanger Sequencing, was performed in individual samples of aforementioned two groups, in the 1^st^ SCLC cohort, 2^nd^ SCLC and CG3 (Fig. 1).

**Fig. 1.**
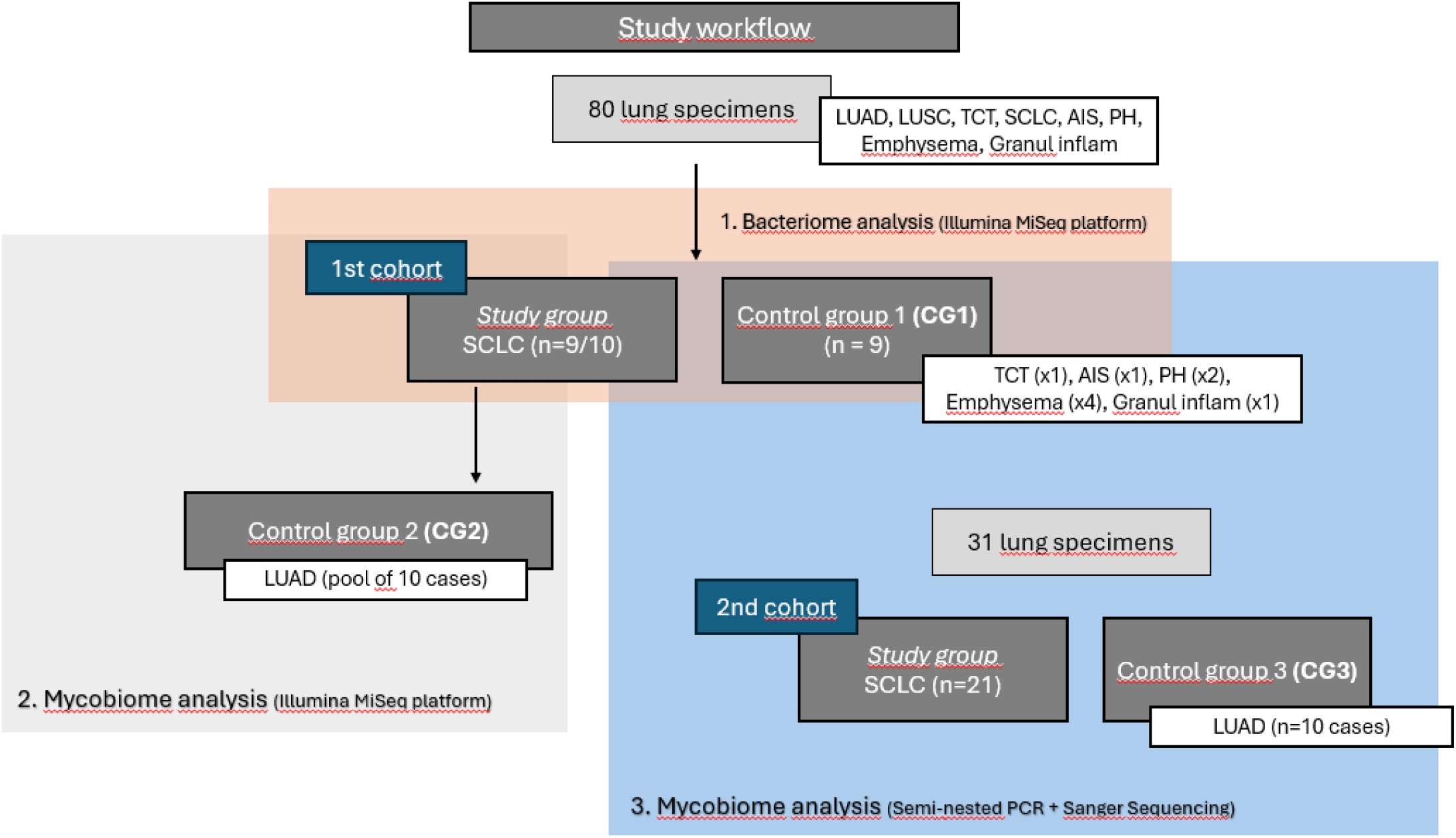
Study Workflow. An initial search of a hospital pathology data base identified 80 lung specimens collected between 2018 to 2020 selected according the histo-pathology criteria to represent eight distinct pulmonary pathologies: lung adenocarcinoma (LUAD), lung squamous cell carcinoma (LUSC), typical carcinoid tumor (TCT), small cell lung cancer (SCLC), adenocarcinoma in situ (AIS), pulmonary hamartoma (PH), emphysema, and granulomatous inflammation - 10 cases per category. Bacteriome Analysis (Illumina MiSeq platform) was performed in the small cell lung cancer (SCLC) samples (1^st^ SCLC cohort, n=10) and a selected control group (CC1, n=9) of samples comprising TCT, AIS, PH (n=2), emphysema (n=4), and granulomatous inflammation 2. Mycobiome Analysis by Illumina MiSeq platform) was studied comparing pools of two groups (1^st^ SCLC cohort (n=10and a control group (CG1, n=10) comprising LUAD samples only. To confirm the mycobiome results a second search was conducted to identify additional cases of SCLC and LUAD, collected between 2012 and 2023 A total of 31 additional cases, corresponding to 21 SCLC and 10 LUAD were selected to enter the study, forming the 2nd SCLC cohort and the third control group (CG3), respectively. Mycobiome Analysis followed a different approach to optimize recovery of fungal DNA sequencing targets: semi-nested PCR followed by Sanger Sequencing.

### Bacteriome Composition

Bacteria identified in paraffin control samples were characterized at the taxonomic level. From the 29 genera shared with the two study groups (1^st^ SCLC and CG1), five genera (*Thermus, Meiothermus, Geobacillus*, Escherichia-*Shigella, Tetragenococcus*) represented 50% of the total bacteria abundance found in the paraffin controls and were subtracted from the microbial datasets of the study groups for *adjustment*, when the ASV ratio (study group/paraffin control) was < 10 arbitrary cutoff.

The Shannon-Wiener index revealed a statistically significant difference in microbial diversity between the 1^st^ SCLC and CG1 (P = 0.003), with lower values observed in the SCLC group (H′ = 2.84) compared to CG1 (H′ = 3.85). This suggests a reduced number of taxa and dominance by a few species in SCLC-associated microbiota. The Effective Number of Species (ENS) further supports this observation, indicating fewer effectively abundant taxa in the SCLC group (ENS = 17) relative to CG1 (ENS = 47). Although the Gini-Simpson Index (1–D), which estimates the probability that two individuals randomly selected from a sample belong to different species, was slightly lower in SCLC (0.91 vs. 0.97), the difference was not statistically significant (P = 0.840). In contrast, Pielou’s Evenness Index (J′), reflecting how evenly individuals are distributed across taxa, differed significantly in the two groups (P = 0.005), with SCLC showing lower evenness (J′ = 0.68) than CG1 (J′ = 0.88) (Fig. 2A, table). Overall, these results demonstrate that the SCLC group harbors a microbiota with significantly lower diversity and evenness, characterized by reduced richness and imbalanced taxonomic distribution. Despite comparable dominance patterns captured by the Gini-Simpson index, the findings point to a disrupted microbial ecosystem in SCLC samples.

**Fig. 2.**
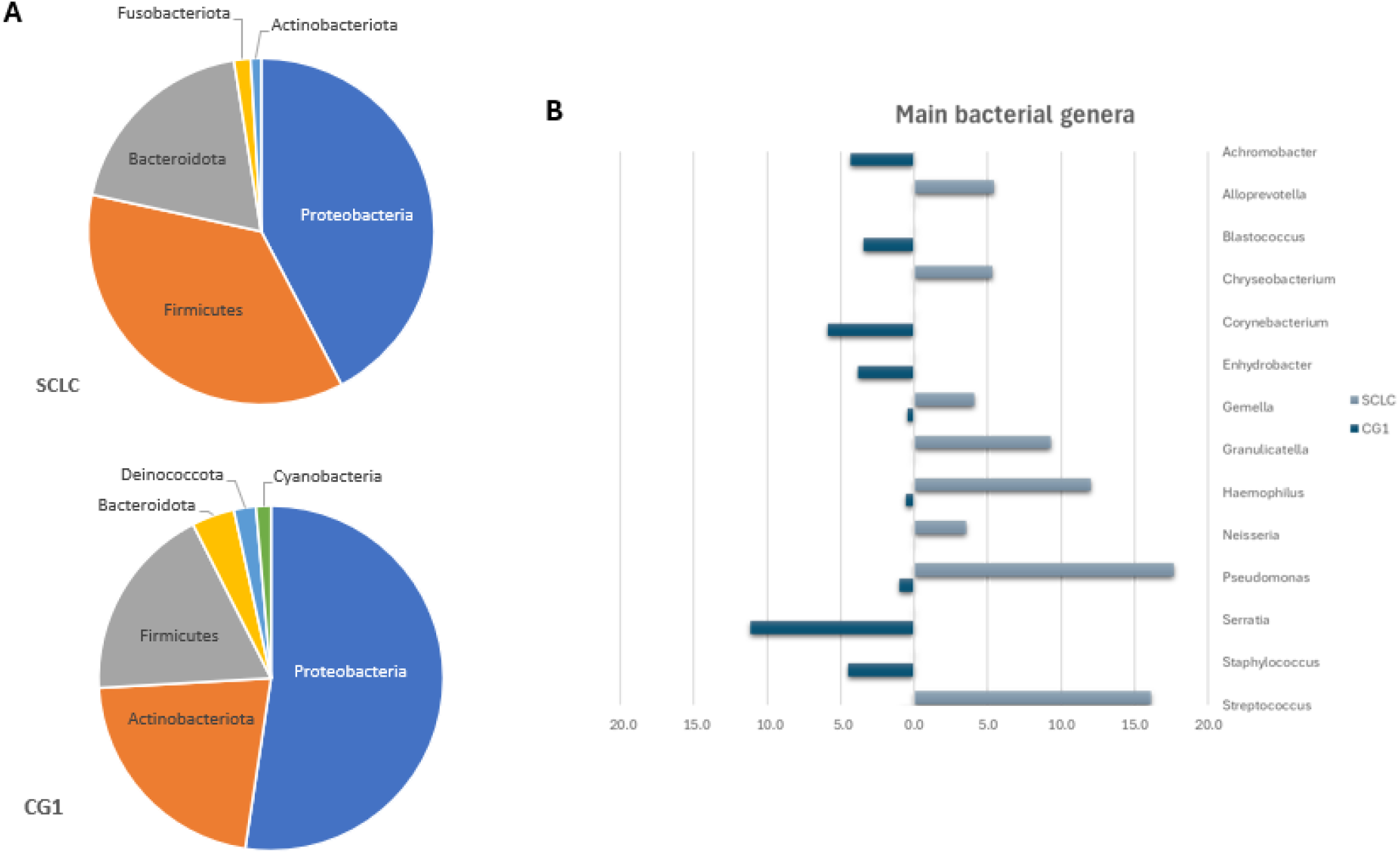
SCLC bacterial diversity. Results from histological samples of SCLC (1^st^ cohort, n=9) and control group 1 (CG1, n=9). **A**. Between groups (alpha) diversity showing reduced diversity in SCLC group as measured by Shannon-Wiener index (H’) distribution in both groups. ***: p = 0.00001, Wilcoxon rank-sum test with continuity correction) (plot) and different alpha-diversity indices. (table). **B**. Within group (Beta) diversity showing increased diversity dispersion in SCLC samples as compared to CG1 samples as illustrated by Principal Coordinate Analysis (PoCA) (plot) and PERMANOVA test (Adonis2 ANOVA) test and Jaccard dissimilarity coefficient (table).

The bacterial β-diversity within the analyzed samples was assessed using Bray-Curtis dissimilarity and visualized through Principal Coordinate Analysis (PCoA) on a two-dimensional plot (Fig. 2B, plot). A PERMANOVA test confirmed that the grouping factor significantly influenced community structure (Pr(>F) = 0.003), indicating a statistically significant difference in bacterial composition within each group. Furthermore, the Jaccard similarity coefficient (J = 0.017) revealed that approximately 98% of taxa were unique to each group, reinforcing that the bacterial community in SCLC samples was distinct from CG1 samples (Fig. 2B, table).

### Bacterial Taxonomy

Bacteria from all samples were characterized at the taxonomic level. In the SCLC group, Proteobacteria and Firmicutes were the most abundant phyla, accounting for 42% and 36% of the community, respectively. In contrast, the CG1 group was dominated by Proteobacteria (52%) and Actinobacteriota (21%), the latter being notably underrepresented in SCLC. Additionally, Bacteroidota constituted a substantial portion of the remaining microbial community in the SCLC group (19%) but was not a dominant phylum in CG1 (Fig.3, pie charts). Interestingly, the Proteobacteria-to-Actinobacteriota ratio was markedly higher in the SCLC group compared to CG1 (43 vs. 2.4), although not statistically significant. *Pseudomonas* (18%), *Streptococcus* (16%), *Haemophilus* (12%), Granulicatella (9%), *Alloprevotella* (5%) and *Chryseobacterium* (5%) were the most abundant genera in SCLC cohort; *Serratia* (11%), *Corynebacterium (6%), Staphylococcus (4%), Achromobacter (4%) and Enhydrobacter (4%)* were the most prevalent genera in CG1 (Fig. 3, bar chart). This analysis shows that the most represented bacterial genera in the two groups are clearly distinct.

**Fig. 3.**
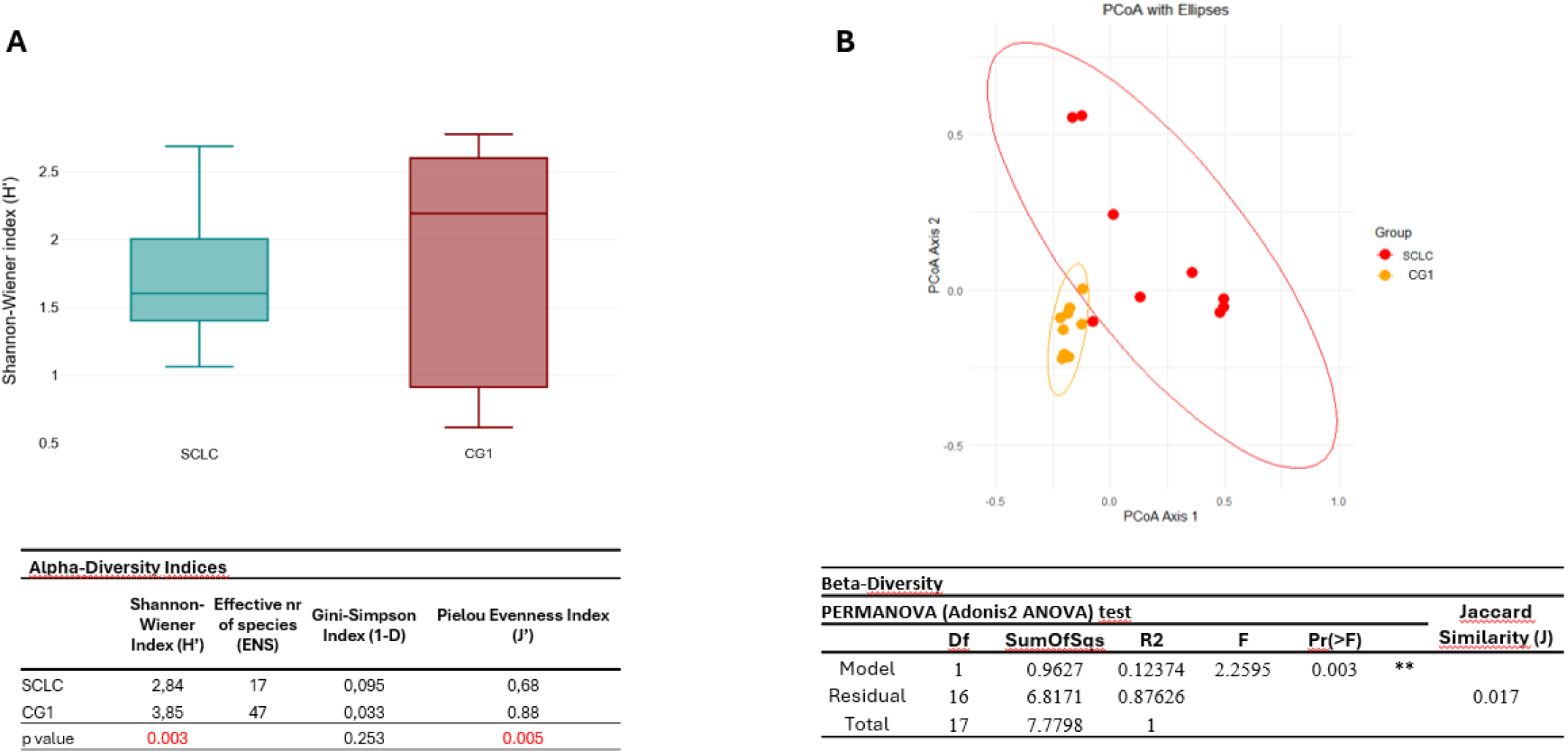
Relative abundance of bacterial taxa. **A**. Main bacterial phyla (pie charts) and **B**. bacterial genera (bar chart) in SCLC and CG1 groups. Relative bacterial abundance >3% for any of the groups in each category were represented.

Overall, the findings highlight a distinct microbial signature associated with SCLC. The SCLC group exhibited a taxonomic profile, characterized by a higher relative abundance of Firmicutes and Bacteroidota, and a markedly different set of dominant genera (*Pseudomonas, Streptococcus* and *Haemophilus*) compared to the CG1, enriched in Actinobacteriota, represented by Corynebacterium.

### Mycobiome Composition

After an initial unsuccessful attempt to analyze each of the ten samples individually, two pooled groups were formed and analyzed: SCLC group (1^st^ SCLC cohort, n=10) and CG2 (LUAD, n=10). *Taphrina* genus, from *Ascomycota* phylum, corresponding to 113 reads, and *Skeletocutis* genus, from *Basidiomycota* phylum, corresponding to 44 reads, were identified in the SCLC group (1^st^ SCLC cohort). *Collembolispora* genus, from *Ascomycota* phylum, was identified residually in CG2 group (3 reads) (Fig. 4A). These striking differences in the SCLC mycobiome composition were compelling enough to explore further.

**Fig. 4.**
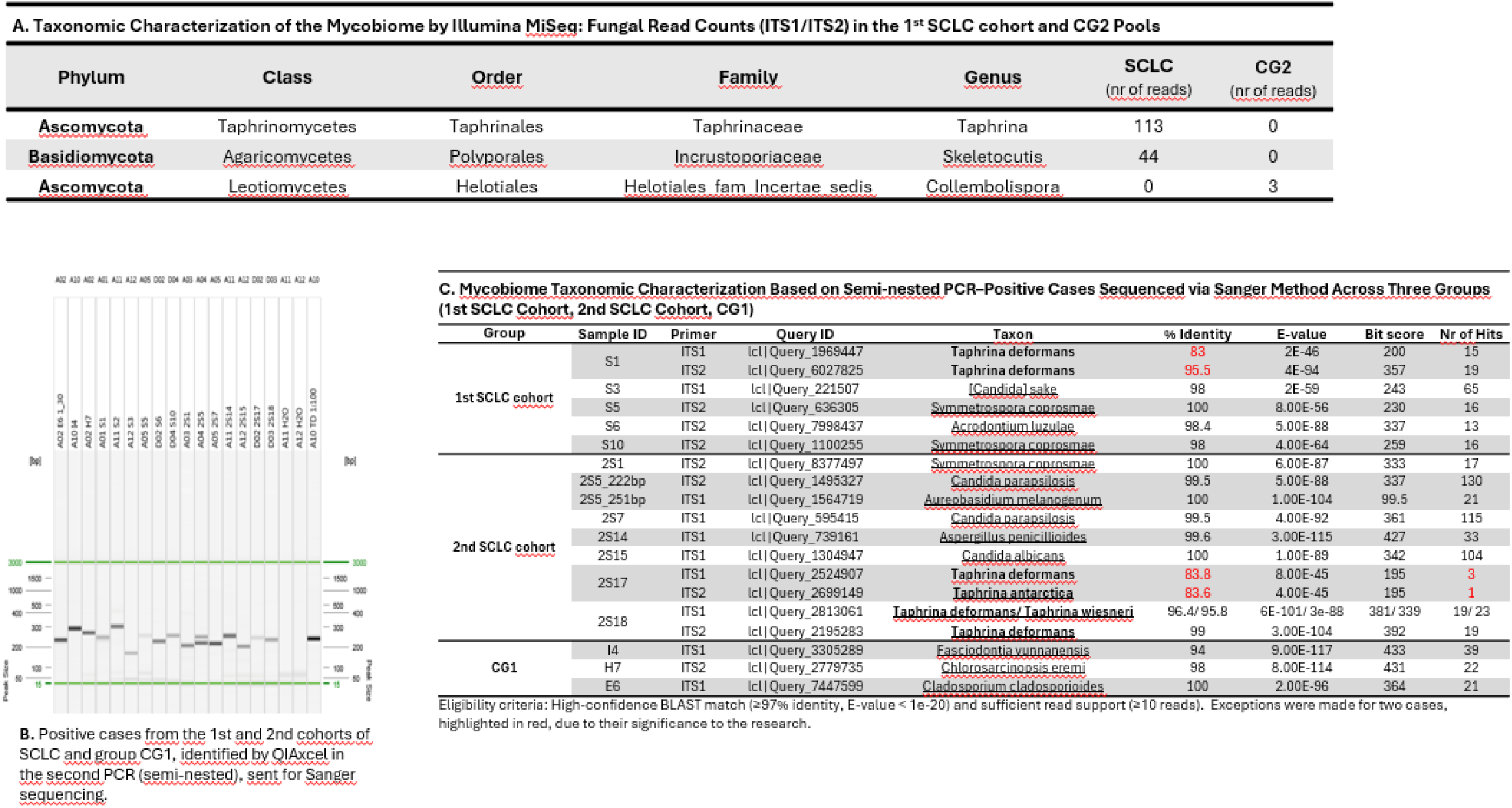
Mycobiome characterization. **A**. Mycobiome taxonomic characterization of Illumina sequencing data: Fungal read counts (ITS1/ITS2) in the 1^st^ SCLC cohort and CG2 Pools. **B**. Gel electrophoresis image of semi-nested PCR products from the 1st and 2nd cohorts of SCLC and group CG1, subjected to Sanger sequencing. **C**. Mycobiome taxonomic characterization of sequenced semi-nested PCR products across three groups (1st SCLC Cohort, 2nd SCLC Cohort, CG1). Cases meeting the adopted eligibility criteria (High-confidence BLAST match (≥97% identity, E-value < 1e-20) and sufficient read support (≥10 reads) are represented. Two exceptions, highlighted in red, were included due to their relevance.

The second sample selection resulted in the addition of the 2^nd^ SCLC cohort (n=21) and the CG3 group (LUAD, n=10). Semi-nested PCR was performed in individual samples of the latter two groups entering the study, as well as on the 1^st^ SCLC cohort (n=9) and CG1 (n=9) groups. Notably, none of the 10 CG3 cases yielded detectable bands on the agarose gel. Bands were detected in 6 of the 9 (67%) cases of the 1^st^ cohort, in 7 of the 21 (33%) cases of the 2^nd^ SCLC cohort and in two (22%) cases of the 9 CG1 group. (Fig. 4B) Sanger sequencing using the ITS1 and ITS2 primers was subsequently performed. Cases meeting the adopted eligibility criteria (High-confidence BLAST match (≥97% identity, low E-value) and sufficient read support (≥10 consistent *Taphrina* sp. reads) were considered for the analysis. In addition, two cases where *Taphrina* sp. were considered: S1 (1^st^ SCLC cohort) and 2S17 (2^nd^ cohort) - both with sequence identity below 97% and the latter also showed <10 reads. Interestingly, *Taphrina deformans* was identified with sequence identities of 83.6%, 95.5% and 96.4% in three (25%) of the twelve eligible SCLC cases. Moreover, two different *Taphrina* species were identified: *T. antarctica* and *T. wiesneri*, with sequence identities of 83.6% and 95.8%, respectively. Detailed taxonomic characterization is summarized in Fig. 4C. These results revealed an unexpected high prevalence of the *Taphrina* genus in SCLC cases.

## DISCUSSION

This study reports, for the first time, the presence of *Ascomycota* phyla, represented by the genus *Taphrina*, in SCLC, identified across two independent cohorts using two distinct methodological approaches. *Taphrina* spp. are known as a plant pathogenic fungi in *Taphrinomycotina* and distinct biological features relating parasitism with dimorphic changes not found in any other plant pathogenic ascomycetes (29). *Taphrina* species generally show strict host specificity, with closely related species infecting only the same or phylogenetically related host species (30). The mechanisms of host adaptation and speciation of pathogenic fungal genomes are diverse, including accumulation of DNA point mutations, chromosomal rearrangement, loss of heterozygosity, ploidy change, and horizontal gene and chromosome transfer (31,32). *Taphrina* species have the smallest known genomes for plant pathogenic fungi, have very low repetitive DNA content and display a noncanonical two-speed genome, with the fast subgenome involved in host adaptation and infection (33). Furthermore, secreted effector proteins can determine the outcome of host-pathogen interaction and host-specificity of pathogenic fungi by modulating host immunity, and many known effector proteins are genus-, species-, or even strain-specific (33). Interestingly, *Taphrina* species are able to produce the plant hormones auxin and cytokinin widely believed to be involved in the many tumor and leaf deformation symptoms caused by various *Taphrina* species (34–36). Fungi have been reported to promote tumorigenesis, however the role and effects of cancer-related fungi are still largely unknown (19,37). A recent study found that most types of fungi have specific bacterial species with which they tend to co-exist, reflecting the close associations between fungi and bacteria (38).

Bacteria in lung cancer are characterized by a decline in α-diversity and an increase in total bacteria load (39). In this study, the SCLC samples exhibited reduced richness and imbalanced taxonomic distribution characterized by dominance of few species. Additionally, these results also revealed a significant difference in β-diversity between the two groups, with SCLC samples showing higher dissimilarity, reinforcing the notion that the microbial profile SCLC has distinctive features. Although findings across published studies vary, *Proteobacteria, Firmicutes*, and *Bacteroidota* consistently emerge as the most frequently identified phyla in healthy lung microbiota. At the genus level, *Pseudomonas, Streptococcus, Prevotella, Fusobacteria* and *Veillonella* predominate, with lesser contributions from potential pathogens including *Haemophilus* and *Neisseria* (40). A similar abundance distribution at phyla and genera level was demonstrated in the SCLC group: *Proteobacteria, Firmicutes* and *Bacteroidota* were the most abundant phyla. The most abundant genera identified were *Pseudomonas, Streptococcus, Haemophilus, Granulicatella, Alloprevotella*, and *Chryseobacterium*, aligning with previous studies on bronchial brush specimens in lung cancer patients, that report a gradual increase in *Streptococcus* and a decline in *Staphylococcus* along the continuum from healthy tissue to non-cancerous lesions and ultimately to cancerous sites (41). Moreover, the relative abundance of Actinobacteria in SCLC group is decreased, corroborating one of the findings of the meta-analysis on lung microbiome and lung cancer by Najafi et al. (42). Furthermore, there is an increase in the Proteobacteria-to-Actinobacteria ratio in tumoral tissue - an indicator of microbial balance or dysbiosis in gut microbiome profiling from fecal samples - with higher ratios suggesting a shift toward a more inflammatory gut environment (43–45). As far as we are aware, no other study reported on SCLC microbiota. The taxonomic analysis uncovered distinct compositional profiles between the two groups, highlighting a divergent microbial landscape in SCLC that may reflect - or potentially contribute to - disease-related alterations in community structure. A key limitation of this analysis is that the samples were not initially collected for microbiome analysis. This study has taken steps forward by describing for the first time the microbiome of SCLC, a low frequency lung tumor. Our study unveiled a potential association of the biotrophic plant pathogenic fungi genus *Taphrina* with human cancer, raising the possibility that a new adaptation of this strictly host-specific known-plant pathogen should underly the colonization of human lung tumors, namely SCLC. While this finding is indeed fascinating, its role as a casual bystander or hidden culprit is unclear. Nevertheless, the involvement of *Taphrina* spp. and their metabolic products in specific pathogenic processes of this type of cancer warrants thorough investigation.

## Acknowledgements

The authors gratefully acknowledge Professor Jason Slot from the Department of Plant Pathology at The Ohio State University for his insightful comments, constructive suggestions, and generous support. The authors thank the Interventional Pneumology, Thoracic Surgery, and Interventional Radiology teams at the Champalimaud Clinical Centre for invaluable collaboration in sample collection.

